# Multi-omics profiling links epigenetic and lncRNA changes to early human endochondral ossification priming

**DOI:** 10.64898/2026.07.08.735777

**Authors:** David Hidalgo Gil, Alejandro Garcia Garcia, Francine Wolf, Sara González Antón, Sacha Bosch, Ani Grigoryan, Phuong Cao Thi Ngoc, Sowndarya Muthukumar, Cristian Bellodi, Andrea Barbero, Paul Bourgine

## Abstract

Differentiation programs remain incompletely understood across stem cell types, including for human bone marrow mesenchymal stromal/stem (BM-MSCs) cells, a heterogenous population orchestrating bone formation and establishing a functional hematopoietic niche in the bone marrow. BM-MSCs form and repair bone through the evolutionarily conserved process of endochondral ossification (EO), initiated by deposition of a transient cartilage template subsequently remodeled into bone and bone marrow tissues. Despite their considerable potential for skeletal regeneration, the early molecular and cellular events underlying BM-MSCs commitment to EO remain elusive. To overcome donor dependent variability in chondrogenic potential that limits mechanistic studies, we here exploit OssiGel as a potent chondro-inductive extracellular matrix offering robust recapitulation of EO by BM-MSCs. Through multi-omics profiling of OssiGel-primed BM-MSCs, we identify rapid chromatin remodeling at chondrogenic loci as concomitant with lineage commitment. The emergence of chondro-progenitors and TWIST1+ skeletal populations is detected as early as 3 days *in vivo* and correlates with successful EO recapitulation. Mechanistically, we identify LINC02511 as positive regulator of EO. We confirm presence of LINC02511 in human skeletal atlases, and its CRISPR-mediated silencing significantly impaired EO. By integrating human tissue engineering strategies with single cell multi-omics profiling, our study provides a framework for deciphering BM-MSCs fate decisions, highlighting the role of enhancer and non-coding elements as key determinants of early lineage specification. These findings advance our understanding of BM-MSCs biology and will prompt their translational exploitation in regenerative medicine.

## Introduction

Endochondral ossification (EO) is the principal developmental program governing bone formation and repair of the axial skeleton. During embryogenesis, this process is initiated by the proliferation and aggregation of a mesodermal-derived population defined as Mesenchymal Stem Cells (MSCs). Early activation of the master transcription factor Sry-Box 9 (SOX9)^1^ primes MSC differentiation into proliferative chondrocytes, leading to cartilage formation. Signaling gradients of fibroblast growth factors (FGFs), bone morphogenic proteins (BMPs) and parathyroid hormone related protein (PTHrP) further orchestrate chondrocyte maturation into hypertrophic chondrocytes, marking the onset of the cartilage-to-bone transition^2^. Concomitantly, a subset of hypertrophic chondrocytes begins expressing Runt-related transcription factor 2 (RUNX2) and Osterix, driving osteoblast emergence^3^ and initiating ossification. Terminally differentiated osteoblast can subsequently mature into osteocytes^4^, which become embedded in the mature bone matrix^2^. Upon completion of this ƒleaprocess, a fraction of MSCs persist within the established bone marrow (BM), forming the BM-MSCs population present in adulthood and maintaining intrinsic skeletal differentiation capacity.

In humans, the genetic and epigenetic events governing BM-MSCs differentiation during the EO process remain incompletely understood. Advances in epigenomic profiling have revealed lineage specific chromatin landscape underlying the osteogenic and adipogenic differentiation of BM-MSCs^5^. However, a similar mechanism linked to chondrogenic priming -and thus EO initiation-remains unknown. Studies have identified specific chromatin accessibility patterns in BM-MSCs^6^, correlating with enhanced EO capacity. Chromatin immunoprecipitation analysis^7^ further suggested an “endochondral signature” linked to a BM-MSC-specific enhancer landscape. Fundamentally, these findings established a first connection between epigenetic status and differentiation potential of BM-MSC populations, distinct from that of extra-skeletal MSC sources.

Despite recent developments, several critical gaps remain. Existing studies are largely limited to bulk populations, precluding the identification of functionally distinct sub-populations within the BM-MSCs compartment. Moreover, comparisons have typically been restricted to undifferentiated or terminal cell states, failing to capture the early molecular events that trigger EO. This raises the question of whether the EO potential of BM-MSCs is imprinted in genome accessibility patterns, whether mesenchymal populations homogenously respond and drive chondrogenesis and if early regulatory mechanisms controlling differentiation can be identified. BM-MSCs represents the leading cell source in regenerative medicine ^8–11^, addressing these questions is essential from both a fundamental standpoint and towards increased translation of BM-MSCs therapies.

Building on advances in tissue engineering and principles from developmental biology^12,13^ a strategy has been proposed to recapitulate human EO. This “developmental engineering” approach relies on a two-step protocol, starting by the *in vitro* chondrogenic priming of BM-MSCs, cultured either as micromass pellet ^14^ or within collagen scaffolds^15^. Following 3 to 6 weeks this enables mature human hypertrophic cartilage formation. In a second step, ectopic implantation of generated cartilage templates into immuno-compromised mice recapitulates the latest stages of EO, ultimately generating a mature bone organ with persisting BM-MSC populations^16^. While variants of this protocol have been reported ^17,18^, a common limitation remains the highly variable BM-MSCs chondrogenic differentiation *in vitro*, a pre-requisite for effective EO *in vivo*. This yet-to-explain donor-to-donor variability combined with the lack of performance predictive markers ^19–21^ challenges both the translational exploitation of BM-MSCs and the mechanistic study of EO.

Here, we propose exploiting OssiGel™, a human-derived extracellular matrix with chondro- and osteo-inductive properties to enable more reproducible EO priming^22^. By combining BM-MSCs with OssiGel, we aim at uncovering the early molecular and epigenetic determinants of BM-MSCs commitment towards EO using a multi-omics approach at single-cell resolution. Our findings reveal the emergence of a TWIST1+ mesenchymal skeletal stem cell and chondro-progenitor populations as early as 3 days after EO priming. This transition is associated with chromatin remodeling at early stages of EO, while BM-MSCs show a pre-defined osteogenic chromatin landscape^5^ which is further tailored in specific cell types. We further report the identification of the long noncoding RNA LINC02511 as a novel regulator of EO, functionally validated as an enhancer element using CRISPR-Cas9 silencing and *in vivo* assays.

In summary, our work contributes to decoding the human regulatory processes governing stem cell fate decisions exemplified in the EO context. From a translational perspective, it can prompt the exploitation of BM-MSC populations for skeletal repair^21^, where EO approaches remain to be clinically implemented.

## Results

### OssiGel confers superior chondrogenesis priming of human bone marrow mesenchymal stem/stromal cells and robust recapitulation of endochondral ossification

OssiGel is an engineered human BM-MSC extracellular matrix (Fig. S1A) with previously demonstrated osteoinductive properties^22^. We first aimed at evaluating its capacity to support BM-MSCs chondrogenesis *in vitro* across multiple donors. To this end, BM-MSCs were isolated from healthy BM samples and *in vitro* expanded prior to differentiation. BM-MSC samples were initially stratified based on their chondrogenic capacity (CC) pre-determined by modified Bern score (MBS) ^23,24^ after chondrogenic differentiation in collagen sponges. Highly chondrogenic donors (High-CC) were defined with a score superior to 3, while donors with low chondrogenic capacity (Low-CC) had a score below 3. Next, three *in vitro* culture conditions were compared, consisting in cell seeding on collagen sponge (CSP) as a gold-standard procedure^15^, embedding in Fibrin/thrombin hydrogel (Fibrin), or embedding in OssiGel (Fig. 1A). During 3D culture, cells were exposed to standard chondrogenic medium for 3 weeks.

**Figure 1.**
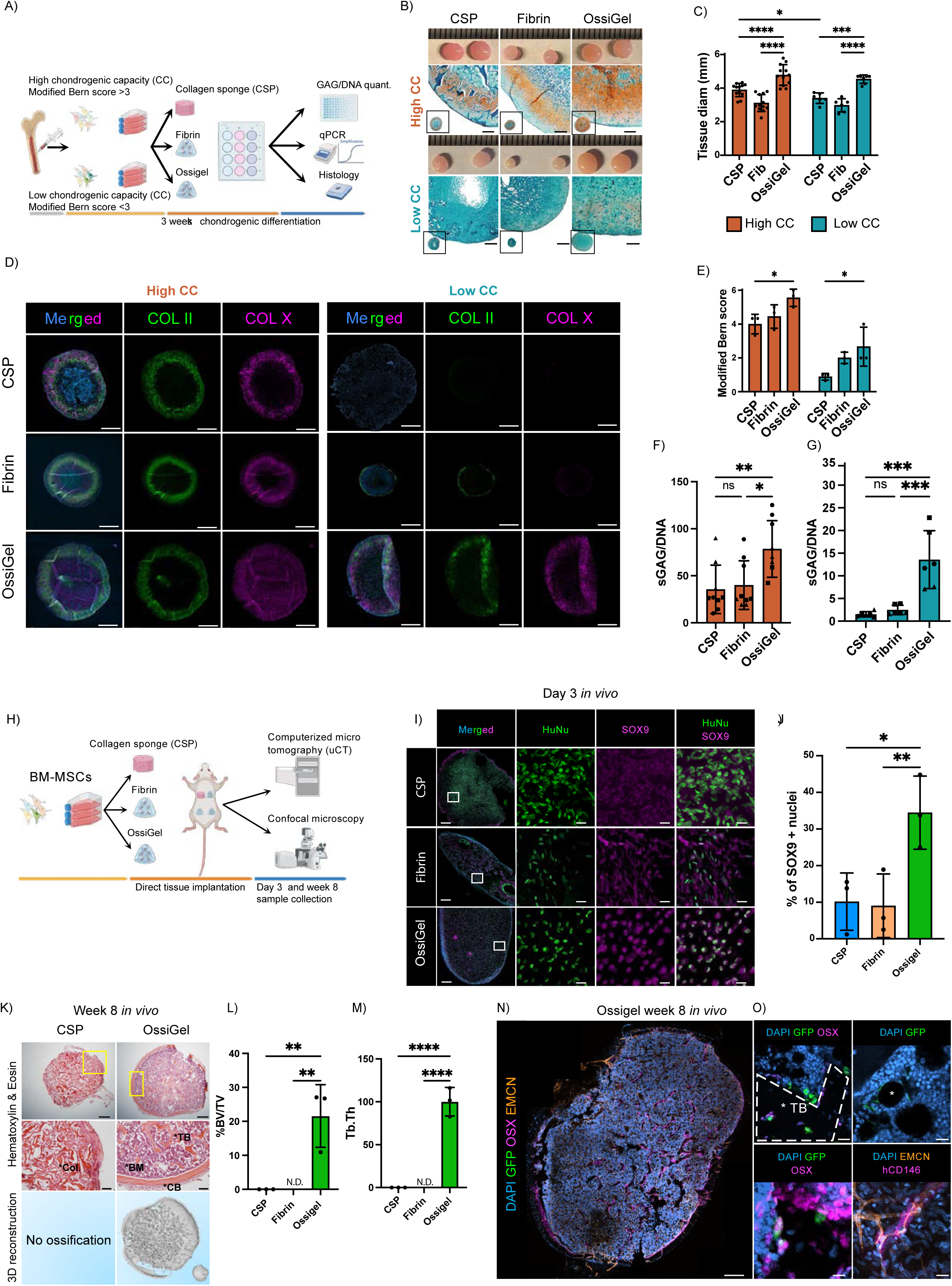
OssiGel confers superior chondrogenesis priming of human bone marrow mesenchymal stem/stromal cells and robust recapitulation of endochondral ossification. (A) Schematic of the experimental workflow for isolation, chondrogenic differentiation, and analysis of primary BM-MSCs into cartilage tissues. (B) Representative macroscopic images and Safranin-O staining after 3 weeks of chondrogenic differentiation in CSP, fibrin, or OssiGel scaffolds, from donors with high (top) or low (bottom) chondrogenic capacity (CC). Scale bars, 500 µm. (C) Tissue diameter by scaffold condition grouped by CC after 3 weeks of chondrogenic differentiation. (D) Representative immunofluorescence for type II and type X collagens in high-CC (left) and low-CC (right) samples. Scale bars, 1 mm. (E) Modified Bern scores of engineered cartilage from primary BM-MSCs cultured in CSP, Fibrin and OssiGel grouped by CC. (F and G) Sulfated glycosaminoglycan (sGAG)–to–DNA ratio after 3 weeks of chondrogenic differentiation in CSP, fibrin, or OssiGel scaffolds for high-CC (F) and low-CC (G) samples. (H) Schematic of the experimental design for *in vitro* construct formation and subcutaneous implantation in NSG mice, analyzed at 3 days, 6 weeks and 8 weeks post-implantation. (I) Immunofluorescence of 3-day implants for DAPI, human nuclei (HuNu), and *SOX9*. Scale bars, 500 μm (overview) and 15 μm (magnified regions). (J) Quantification of SOX9-positive nuclei in CSP, fibrin, and OssiGel conditions at 3 days. (K) Hematoxylin and eosin staining of 8-week explants showing unremodelled collagen sponge (Col.), cortical bone (CB), bone marrow (BM), and trabecular bone (TB). Scale bars, 500 μm (overview)and 100 μm (magnified regions) and µCT reconstruction of implants at 8 weeks *in vivo* (bottom panel). (L) Micro-computed tomography (μCT) quantification of bone volume/total volume (BV/TV) in recovered CSP tissues and OssiGel-derived ossicles at 8 weeks. (M) Trabecular thickness quantification in recovered CSP and OssiGel derived ossicles at 8 weeks. (N) Immunofluorescence of 8-week OssiGel ossicles for GFP-labeled BM-MSCs, osterix (OSX), and endomucin (EMCN), showing osteogenic differentiation and vascularization. Scale bar, 1 mm. (O) Magnified regions identifying stromal lineages: adipocytes (top left), osteocytes (top right), osteoblasts (bottom left), and perivascular stroma (bottom right). Data represents means ± SD; n = 6 independent BM-MSC samples, 3 per group. *P < 0.05, **P < 0.01, ***P < 0.001 by one-way ANOVA (F, G, J, L, M) or two-way ANOVA (C, E).

Following 3D differentiation, OssiGel-derived tissues appeared significantly larger compared to CSP and Fibrin tissues (Fig. 1B, 1C). Safranin-O staining of High-CC donors confirmed deposition of a sulfated glycosaminoglycans (sGAG) rich extracellular matrix in all conditions, however substantially increased in the OssiGel group (Fig. 1B). Low-CC donors consistently showed poor cartilage deposition, although the increased tissue size in the OssiGel group was associated with detectable cartilage areas and chondrocyte-like cell morphology (Fig. 1B, Fig. S1B). Immunofluorescence staining was further performed to assess the presence of Collagen type-II and Collagen type-X, hallmarks of hypertrophic cartilage. The OssiGel group exhibited the strongest collagen deposition from both High-CC and Low-CC donors (Fig. 1D, Fig. S1C). Engineered tissue quality was further assessed quantitatively, first using Bern scoring which revealed that OssiGel consistently increased the cartilage formation across donors as compared to Fibrin and CSP conditions (Fig. 1E). In line with this finding, OssiGel also resulted in the highest sGAG/DNA ratio with a 2-fold and 4-fold increase as compared to the CSP condition in High-CC (Fig. 1F) and Low-CC samples (Fig. 1G), respectively. Interestingly, DNA quantification showed no significant differences across conditions, excluding proliferative effects as an explanation for increased sGAG accumulation in OssiGel cartilage tissues (Fig. S1D-E). Collectively, this establishes OssiGel as a potent chondro-inductive matrix superior to standard 3D in vitro protocols and validated across multiple BM-MSC donors.

Conventional protocols require *in vitro* cartilage formation followed by implantation into animal models to recapitulate EO^14^. Here, the OssiGel technology was further evaluated to determine whether exposure of BM-MSCs to OssiGel followed by direct ectopic implantation is sufficient to induce EO and generate mature bone and BM *in vivo*. To this end, BM-MSCs (three independent donors per group) were seeded onto CSP, embedded in either fibrin or OssiGel, and implanted into immunocompromised mice without prior *in vitro* differentiation (Fig. 1H, Fig. S1F). Tissues were isolated 3 days, 6 weeks and 8 weeks post-implantation to characterize early and late stages of EO development.

At 3 days post-implantation, macroscopic examination revealed greater vascularization around OssiGel than around CSP and fibrin implants (Fig. S1G). Immunofluorescence staining showed SOX9 - a master transcription factor of the chondrogenic lineage - detectable in all conditions (Fig. 1I). However, SOX9 was predominantly nuclear in the OssiGel and mostly cytosolic in the CSP and Fibrin groups (Fig. 1I–J), suggesting greater SOX9 activation and initiation of the endochondral program in OssiGel implants. Histological analysis of OssiGel tissues explanted after 6 weeks confirmed successful initiation of EO, evidenced by presence of cartilage, cortical bone and an emerging bone marrow cavity (Fig. S1H).

The remaining tissues were explanted 8 weeks after implantation to assess EO outcomes in each condition. OssiGel- and CSP-derived tissues were consistently retrieved, whereas the Fibrin constructs had been fully resorbed (Fig. S1I). CSP samples consisted of fibrous tissues, with no histological evidence of bone formation or mineral deposition by μCT (Fig. 1K, Fig. S1I-J). In sharp contrast, OssiGel constructs fully remodeled into bone organs (Fig. 1K), with confirmed presence of cortical and trabecular structures, and a central bone marrow compartment (Fig. 1K, Fig. S1I-J). μCT reconstructions further revealed their 3D bone architecture and enabled quantification of bone volume relative to total tissue volume (BV/TV, Fig. 1L) and trabecular thickness (Fig. 1M). These features were consistently observed in OssiGel tissues and absent from the other samples.

Finally, the fate of implanted human BM-MSCs and their contribution to tissue formation were assessed by immunofluorescence analysis. Before implantation, human BM-MSCs were GFP-labelled by lentiviral transduction, enabling cell tracking alongside staining with human-specific antibodies (Fig. 1N-O, Fig. S1K). In OssiGel tissues, human BM-MSC-derived stromal cells, CD146⁺ perivascular stromal cells, and adipocytes with large lipid vacuoles were identified in the marrow compartment (Fig. 1O). Bone-associated human cells were also present: Osterix (OSX) staining indicated osteoblasts lining bone structures and osteocytes embedded within the bone matrix (Fig. 1O). These validate the presence of multiple mesenchymal lineages in OssiGel tissues following EO priming. By contrast, no human cells were detected in CSP tissues, indicating a lack of cell persistence and differentiation in this condition (Fig. S1K).

Taken together, these findings demonstrate that OssiGel-primed BM-MSCs can effectively recapitulate EO *in vivo* without prior *in vitro* chondrogenic differentiation, producing remodeled tissues containing bone and bone marrow. Notably, BM-MSCs also give rise to canonical mesenchymal lineages, contributing to the establishment of a human-like bone marrow niche.

### Early emergence of Chondro-progenitors and TWIST1*+* mesenchymal skeletal stem cells is a hallmark of endochondral ossification priming

OssiGel reproducibly drives endochondral priming of BM-MSCs across donors, allowing the study of early molecular events of the human EO pathway. To this end, single-nucleus multiomic profiling (RNA-seq and ATAC-seq) was performed on healthy donor BM-MSCs before and after endochondral priming. BM-MSCs were expanded in vitro and divided into three groups: one was profiled directly to establish baseline RNA-expression and chromatin-accessibility profiles, while the other two were embedded in OssiGel or Fibrin and implanted in vivo under identical conditions (Fig. 2A). Fibrin served as a negative control because it did not support EO. After 3 days, human nuclei were retrieved from the implants using a nattokinase-based protocol^25^. Following quality filtering, an RNA-seq and ATAC-seq dataset composed of 15,650 nuclei was generated (three independent BM-MSC donors).

**Figure 2.**
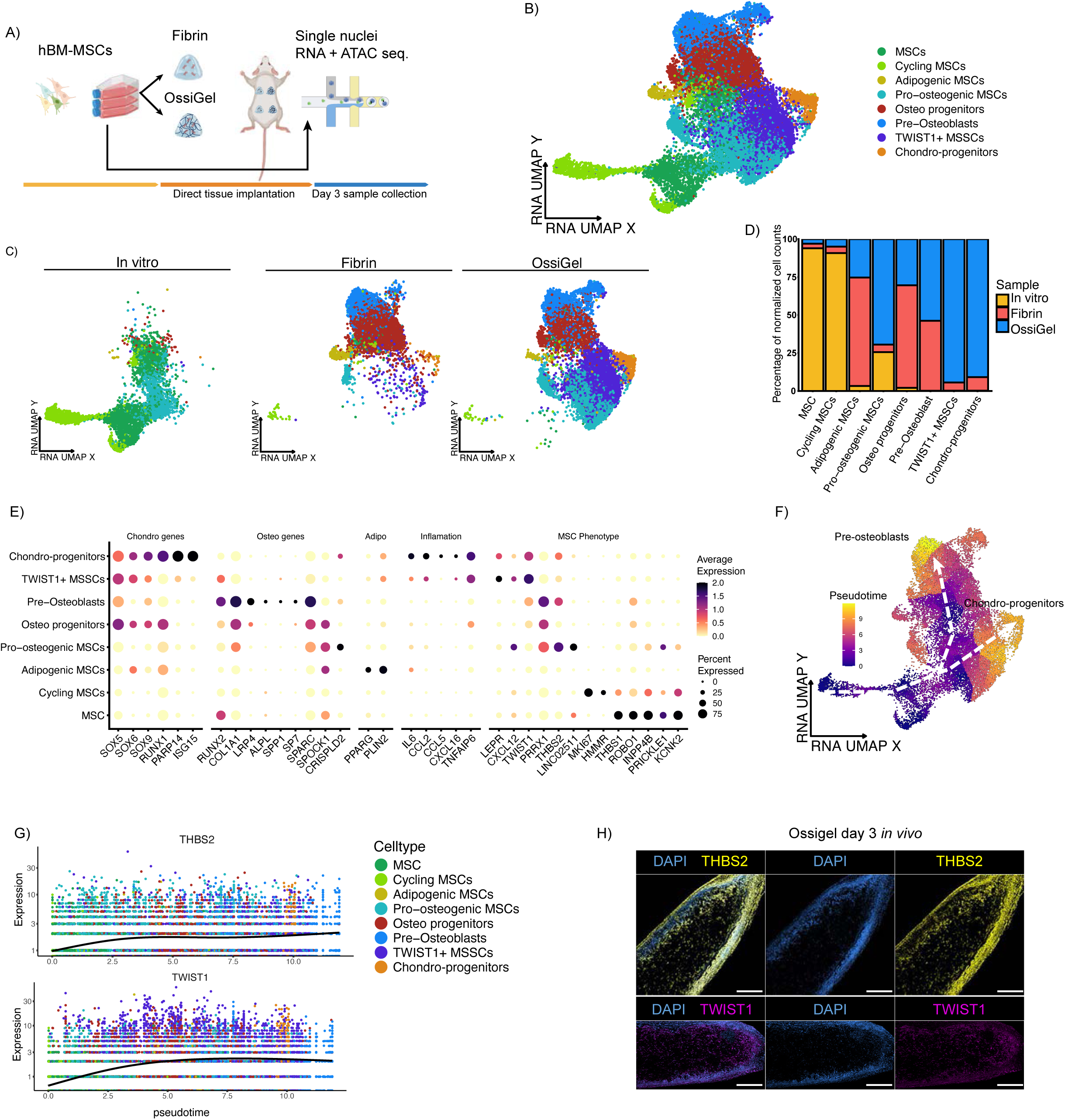
Early emergence of Chondro-progenitors and TWIST1*+* mesenchymal skeletal stem cells is a hallmark of endochondral ossification priming. (A) Schematic of the experimental design for *in vitro* construct formation and subcutaneous implantation in NSG mice, and single nuclei ATAC + RNA seq after 3 days *in vivo*. (B) UMAP embedding of single-nuclei RNA-seq data at 3 days post-implantation. (C) UMAP embedding of single-nuclei RNA-seq data split by condition, color represents cell cluster. (D) Bar plot of cluster composition from each respective sample. (E) Dot plot of average scaled expression (color) and fraction of expressing cells (dot size) for selected marker genes. (F) Pseudotime trajectory on UMAP embedding. (G) Pseudotime ordered expression of THBS2 and TWIST1. (H) Immunofluorescence of 3-day *in vivo* OssiGel for THBS2 and TWIST1. Scale bar 500µm.

Initial clustering based on gene expression analysis identified 8 distinct populations with distinct distribution across samples (Fig. 2B-D). Of note, at this early timepoint, cells maintained the expression of canonical MSC markers (Fig. S2B) although critical differences in expression level and marker types were present across clusters. The In vitro group was predominantly composed of the MSCs and Cycling MSCs clusters, defined as the less committed populations. The MSCs cluster (Fig. 2C) was primarily characterized by a strong expression of *PRICKLE1, ROBO1* -an osteogenic inhibitor^26^, *THBS1* and *INPP4B* involved the PI3K/Akt signaling pathway which control cell growth and survival. Interestingly, this cluster also display a relatively high *RUNX2* expression, suggesting pre-activation of the osteogenic lineage. The Cycling MSCs population expressed *MKI67* as key regulator of cell cycle related genes, and the *HMMR* receptor involved in cell migration and repression of adipocytic differentiation^27–30^ (Fig. 2E). Interestingly, the *in vitro* condition also exhibited a fraction of the Pro-osteogenic MSCs cluster (Fig. 2C), a population massively present in the OssiGel group. This cluster displayed high expression of *COL1A1*, *SPARC* and *SPOCK1*, together with the skeletal progenitor markers *PRRX1* and *THBS2* (Fig. 2E).

Following *in vivo* implantation, more committed lineages were primarily identified in both Fibrin and OssiGel conditions. Among those, the Adipogenic MSCs cluster, limited in size, displayed high expression levels of *SOX6*, *PPARG* and *PLIN2*^31^. The *in vivo* groups also shared the Osteo progenitors cluster, characterized by higher levels of *COL1A1*, *SPARC* (encoding osteonectin) and *SPOCK* (testican-1) as well as exhibiting lower expression of *SP7* and SOX genes. Following the same trend, we identified a clearly committed Pre-osteoblasts cluster which displayed high expression levels of *RUNX2*, *COL1A1*, as well as the highest expression of *PRRX1*. Their detectable but low expression of *ALPL*, *SPP1*, *SP7* and *SPARC* suggest that those cells are primed but not fully differentiated osteoblasts.

In turn, OssiGel-specific clusters were identified with the TWIST1*+* mesenchymal skeletal stem cells (MSSCs) and Chondro-progenitors populations. TWIST1*+* MSSCs uniquely expressed *TWIST1*, reported to be involved in embryonic bone formation^32^. This cluster also displayed marked expression of *SOX5-6-9*, *RUNX2*, *LEPR^+^* and *CXCL12^+^*, a feature shared with reported bone marrow CXCL12-abundant reticular cells (CAR cells)^33^. Finally, the specific Chondro-progenitors cluster displayed the highest expression levels of SOX5-6-9 genes, as well as *RUNX1*, *PARP14* and *ISG15* described as required for chondrogenic differentiation^34^. Interestingly, this population exhibited an increased expression of inflammatory genes (*IL6*, *CCL2*, *TNFAIP6*) and a high *VEGFA* expression (Fig. 2E, Fig. S2C). This correlates with the previously described increased early vascularization pattern observed at OssiGel implantation sites (Fig. S1G).

To facilitate visualization of cell-state distributions across conditions, we quantified the relative abundance of each cluster in the In vitro, Fibrin, and OssiGel groups (Fig. 2D). This confirmed that TWIST1+ MSSC and Chondro-progenitors clusters were detected exclusively in the OssiGel condition, supporting their involvement in the EO primed by OssiGel. Differences in cluster emergence and commitment were also observed by pseudotime analysis, providing an *in silico* view of cell-state progression by inferred transcriptional trajectories (Fig. 2F). MSCs and cycling MSCs occupied the early states, followed by divergence into two branches: one toward Pre-osteoblasts and the other encompassing TWIST1+ MSSCs and Chondro-progenitors. These patterns suggest divergent cell-state trajectories during endochondral priming.

To resolve cell states along this bifurcation, we profiled signature markers across pseudotime: *THBS2* (Pro-osteogenic MSCs), *TWIST1* (TWIST1+ MSSCs), *SOX9/ISG1*5 (Chondro-progenitors), and RUNX2 (Pre-osteoblasts). These dynamics captured the progressive emergence of pro-osteogenic and TWIST1+ MSSCs populations (Fig. 2G) and late-pseudotime induction of the chondrogenic (*SOX9, ISG15*) in Chondro-progenitors, and osteogenic programs (*RUNX2*) in Pre-osteoblasts (Fig. S2D).

The emergence of Osteo progenitors and TWIST1+ MSSCs populations detected by snRNAseq were further investigated by performing immunostainings for THBS2 and TWIST1 in OssiGel-tissues explanted after 3 days. Both were detected at the protein level within tissues, *not* only confirming the presence of these populations in the tissue but also revealing their distribution. THBS2 cells were shown to accumulate at the tissue periphery, while TWIST1+ cells formed a distinct layer immediately beneath them (Fig. 2H).

Next, to determine whether the observed gene expression changes were associated with lineage-specific regulatory programs, we assessed the functional activation of key transcription factors (TFs) based on the expression of their downstream target genes^35^. This analysis corroborated the activation of the adipogenic TFs PPARG and NRF1 in Adipogenic MSCs. In contrast, activation of the osteogenic TFs RUNX2 and SP7 was restricted to the Pre-osteoblasts cluster, whereas chondrogenic TFs, including SOX5, SOX6, LEF1, and KLF15, were specifically activated in TWIST1+ MSSCs and Chondro-progenitors (Fig. S2E).

In summary, we here report that successful EO priming and recapitulation in OssiGel correlates with the emergence of TWIST1+ MSSCs and Chondro-progenitors as early as 3 days post-priming. Pseudotime analysis revealed progressive activation of chondrogenic pathways, with *SOX5/6/9* and related transcriptional programs preceding subsequent osteogenic commitment marked by *RUNX2* activation.

### Endochondral ossification is associated with an early and distinct chondrogenic chromatin remodeling signatures

Our single nuclei transcriptomic dataset allowed picturing the emergence of phenotypes after EO priming. To determine whether these transcriptional changes were accompanied by alterations in chromatin accessibility, we next profiled the chromatin landscape using snATAC-seq.

Initial clustering of the snATAC-seq data recapitulated the cell types identified in the RNA-seq data and revealed similar cell distributions across conditions (Fig. 3A, B). To assess the contribution of each sequencing modality to the detection of phenotypic changes in BM-MSCs, we compared clustering results obtained from RNA-seq, ATAC-seq, and integrated multiomic data (Fig. S3A). RNA-based analysis revealed fragmented MSC clusters, substantial overlap with more committed cell populations, and limited separation between conditions. For instance, ATAC-seq data alone provided insufficient resolution to reliably distinguish Chondro-progenitors from Adipogenic MSCs using conventional clustering approaches. In contrast, joint embedding of both modalities markedly improved cluster definition and condition separation, resulting in a more coherent representation of the cellular landscape.

**Figure 3.**
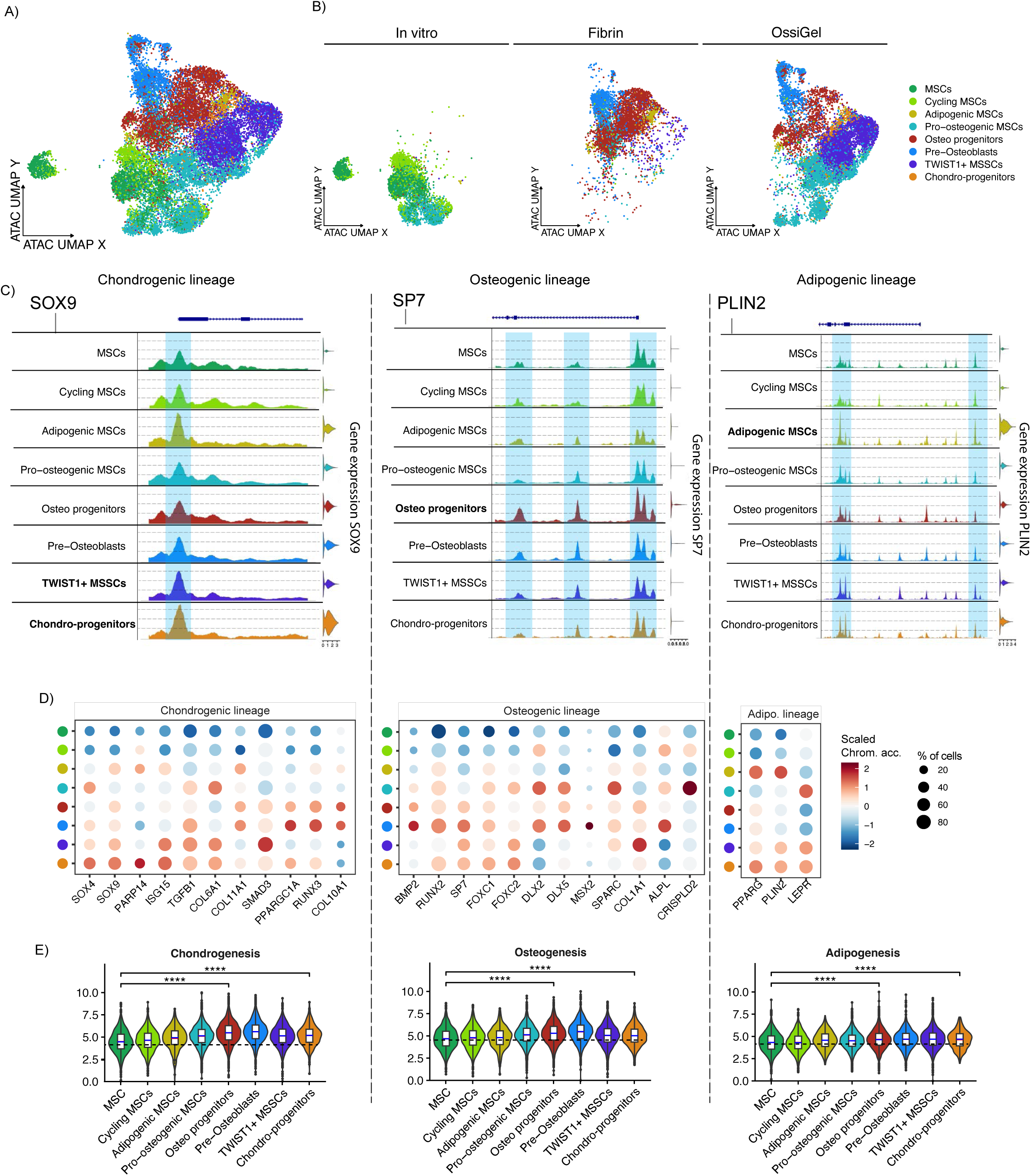
Endochondral ossification is associated with early and distinct chondrogenic chromatin remodeling signatures. (A) UMAP embedding of single-nuclei ATAC-seq data at 3 days post-implantation. (B) UMAP embeddings split by condition; color represents cell cluster. (C) Genome track plots of *SOX9, SP7,* and *PLIN2* loci near promoter regions, with corresponding gene expression levels, highlighted are selected regions of interest. (D) Dot plot of average scaled chromatin accessibility (color) and fraction of cells (dot size) for selected marker genes. (E) Chromatin accessibility scores for lineage-specific genes across cell types.

To first determine chromatin accessibility patterns across the main BM-MSCs lineages, we examined regions associated with master TFs and key marker genes for all identified clusters. This analysis revealed an overall shared baseline of chromatin accessibility at several key gene loci across lineages (Fig. 3C, Fig. S3B), while lineage-specific differences were observed at selected regulatory regions. Regarding the chondrogenic lineage, SOX9 chromatin accessibility at the promoter site doubled in Chondro progenitors and TWIST1+ MSSCs as compared to MSCs, also directly correlating with its higher gene expression. In a similar trend, *PPARGC1A* showed increased accessibility in TWIST1+ MSSCs with higher gene expression as compared to MSCs and Cycling MSCs, clusters restricted to our *in vitro* sample. On the osteogenic lineage, osteogenic genes have been previously described to show moderate chromatin changes after priming^5^. By assessing *SP7* and *ALPL*, we could correlate this observation with minor changes observed at their respective promoter regions. Some differences are nonetheless observed in the case of SP7, particularly in the second exon regions for Osteo progenitors and Pre-osteoblasts clusters. This may indicate a preferential expression of the short version of the SP7 gene^36^ in these populations. Finally, *PLIN2* and *PPARG* genes were selected to account for the adipogenic lineage. Those displayed the most limited remodeling of their loci, despite an increase in corresponding gene expression in Adipogenic MSCs (Fig. 3C, Fig. S3B).

In order to have a quantitative understanding of the chromatin accessibility changes associated with each lineage, we further computed gene activity based on the snATAC dataset. First, a larger gene selection was made to reflect changes in respective chondrogenic, osteogenic and adipogenic lineages (Fig. 3D). This approach confirmed our initial observations with an increased chromatin accessibility in TWIST1+ MSSCs and Chondro progenitors for *SOX9, TGFB1, COL6A1, and SMAD3*. Osteogenic genes such as *BMP2, RUNX2* (also associated with hypertrophy) and *SP7* among others showed increased accessibility confined to the Osteo progenitors and Pre-osteoblasts respectively. Finally, while Adipogenic MSCs showed increased accessibility for *PPARG* and *PLIN2*, the Chondro progenitors cluster displayed a similar accessibility pattern. A second computation strategy was set, by scoring the chromatin accessibility of the gene ontology lists for each lineage (Fig. 3E), thus avoiding biased selection of genes. This method allowed identifying the chondrogenic program as the one associated with the most extensive chromatin remodeling, across all clusters. Changes for the osteogenic were also noticeable, while the adipogenic lineage accounted for the lowest score indicating minor chromatin remodeling.

Next, we compared chromatin accessibility patterns across In vitro, Fibrin and OssiGel conditions (Fig. S3C). This revealed that OssiGel induced the most pronounced increase in chromatin accessibility across all cell clusters and lineages. Notably, increased accessibility at chondrogenic regulatory regions was observed specifically in the OssiGel group, suggesting enhanced commitment toward the EO program. Although less pronounced than in OssiGel, the Fibrin group also exhibited increased chromatin accessibility relative to the In vitro sample, indicating that *in vivo* implantation alone promotes a degree of cellular commitment accompanied by epigenetic reprogramming.

Finally, to validate the activation of differentiation programs, we performed transcription factor motif enrichment analysis based on motif over-representation within accessible chromatin regions. We identified distinct regulatory networks in Chondro progenitors, Osteo progenitors and Adipogenic MSCs clusters (Fig. S3D), characterized by increased activity of TFs associated with their respective differentiation programs, as reflected by enhanced chromatin occupancy.

Collectively, our findings underscore the value of integrated multiomic profiling for resolving early developmental transitions and subtle phenotypic changes in BM-MSCs. We identified a heterogenous and lineage specific BM-MSC response to EO priming, with most significant chromatin access changes occurring in TWIST1+ MSSCs and Chondro progenitors clusters specific to the OssiGel group. These indicate that successful recapitulation of EO is associated with extensive chromatin remodeling and activation of lineage-specific regulatory elements.

### LINC02511 expression is a feature of human embryonic bone development

Our single nuclei transcriptomic dataset further provided us the opportunity to identify novel regulatory elements of the EO program, particularly by comparing OssiGel and Fibrin conditions. We thus performed a differential gene expression analysis allowing the identification of 690 downregulated and 2965 upregulated genes (Fig. 4A). We then focused on non-coding elements since they have been described in some cases to be regulators of cartilage development and osteoarthritis progression ^37–39^. Among a panel of detected long non-coding RNA (lncRNA), *LINC02511* emerged as an interesting candidate regulator due to its marked upregulation in the OssiGel condition (Fig. S4B).

**Figure 4.**
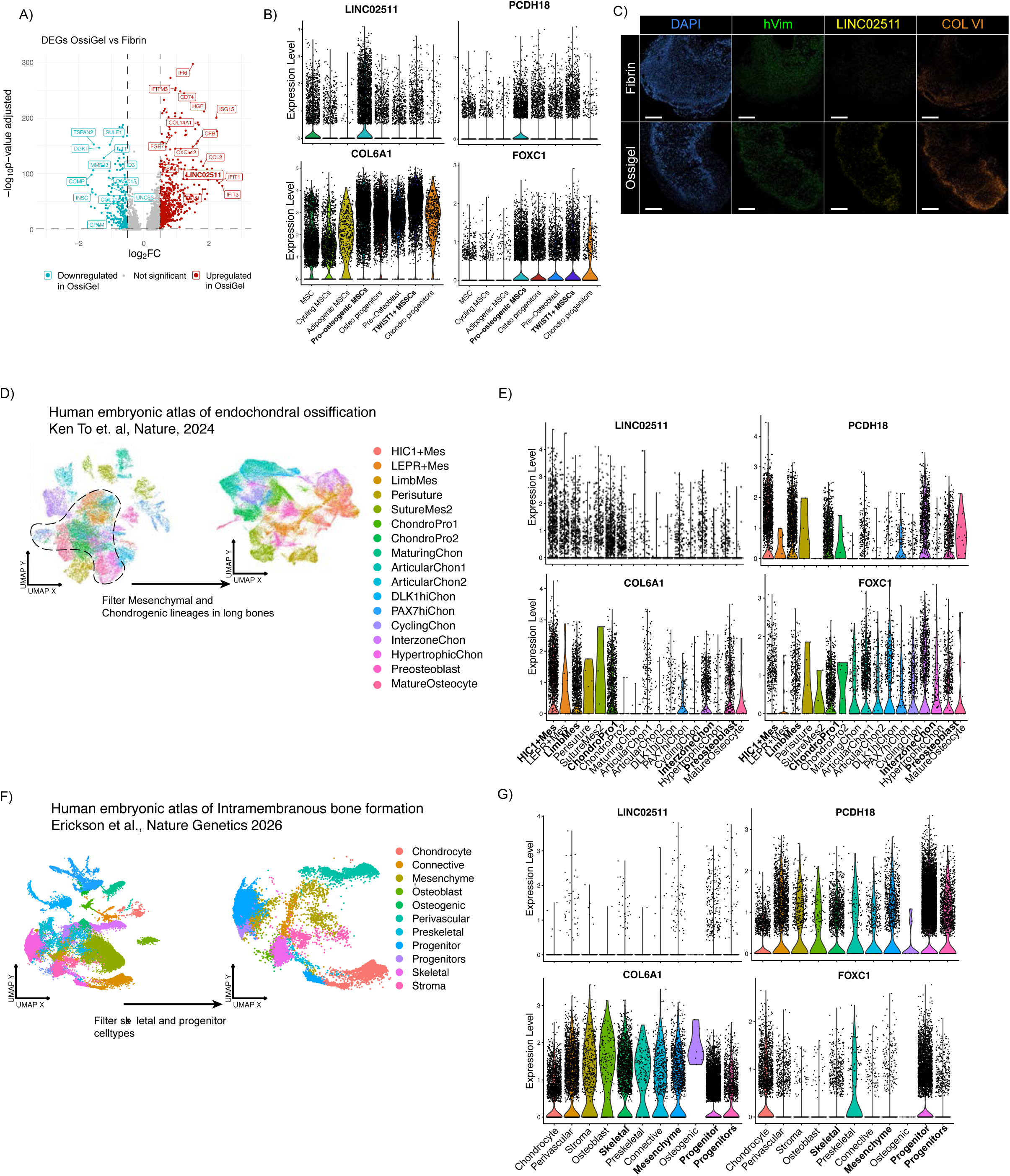
LINC02511 expression is a feature of human embryonic bone development. (A) Volcano plot of differentially expressed genes between OssiGel and fibrin conditions. (B) Violin plot of gene expression of *LINC02511*, and candidate targets *COL6A1*, *PCDH18* and *FOXC1* at 3 days *in vivo.* (C) Immunofluorescence imaging of human vimentin (hVim), COLVI and RNA FISH imaging of LINC02511 in fibrin and OssiGel tissues after 3 days *in vivo*. (D)UMAP embedding of cell types in atlas of human endochondral ossification (left) and of relevant stromal and chondrogenic cell types (right) from (Ken To et. al, Nature, 2024). (E) Violin plot of gene expression of *LINC02511*, and candidate targets *COL6A1*, *PCDH18* and *FOXC1* at across selected cell types in (Ken To et. al, Nature, 2024). (F) UMAP embedding of cell types in atlas of human intramembranous ossification (left) and of relevant stromal and chondrogenic cell types (right) from (Erickson et al., Nature Genetics 2026). (G) Violin plot of gene expression of *LINC02511*, and candidate targets *COL6A1*, *PCDH18* and *FOXC1* at across selected cell types in (Erickson et al., Nature Genetics 2026).

Because lncRNAs can function as epigenetic regulators^38,40^, we *applied in* silico analysis tools to characterize the possible role of *LINC02511* (Fig. S4A). Initial structural analysis performed with RNAfold^41,42^ revealed a complex secondary structure with multiple stem loops, suggesting a possible biological function via physical interactions (Fig. S4C). To investigate this further we applied the LongTarget tool^43^ enabling identification of potential genomic binding sites of LINC02511 on promoter regions across the genome (Supplementary Table 2). Interestingly, top predicted binding targets included *COL6A1, FOXC1* and *BRSK2*. Those are respectively described markers of early chondrogenesis, osteogenesis and regulatory element shown necessary for direct reprogramming of placenta-derived MSCs into chondrocytes. In addition, multiple cartilage-associated collagens including *COL6A1-3, COL5A1, COL18A1*, and *COL20A1* were identified.

We further applied catRAPID v2.1^44^ to predict RNA-protein interactions (Supplementary table 3). Gene ontology enrichment analysis of high-scoring predicted interactions showcased an increased enrichment on gene expression and nucleotide metabolism, compatible with a possible role as an expression enhancer (Fig. S4D). Finally, sequence homology analysis showed *LINC02511* as highly conserved across primates, with partial homology in mouse (Fig. S4E). Phylogenetic conservation analysis across 100 vertebrate genomes revealed low preservation at the LINC02511 locus, but high preservation between primate species (Fig. S4E-F).

To explore the genomic context of *LINC02511*, we analyzed its local syntenic neighborhood, as conserved chromosomal organization frequently marks shared regulatory units. Among the flanking genes, we identified *PCDH18*, which has been linked to robust osteogenesis in bone-forming BM-MSCs ^45^.

Guided by these in silico findings, *LINC02511, PCDH18, COL6A1*, and *FOXC1* expression was examined in our dataset for potential co-expression (Fig. 4B). Populations with high LINC02511 expression also expressed *PCDH18, COL6A1,* and *FOXC1*, particularly pro-osteogenic MSCs and TWIST1⁺ MSSCs. This finding was further examined by RNA FISH for *LINC02511* and immunostaining for collagen VI in Fibrin and OssiGel tissues 3 days after in vivo implantation. LINC02511 was detected only in OssiGel tissues, where its presence coincided with strong collagen VI staining (Fig. 4C). *LINC02511* expression could be linked to what we defined as Pro-osteogenic MSCs in our dataset and co-expressed with *THBS2* key marker in MSC fate and matrix development^46^ displaying a similar tissue distribution.(Fig. 2E,H).

Because *LINC02511* has not previously been linked to BM-MSC differentiation, its expression was examined in two published human embryonic single-cell atlases covering the developmental pathways that form bone: EO (Ken To et al. ^34^) and intramembranous ossification (Erickson et al.^47^). The EO atlas covered several bone-forming regions and included 47 cell clusters. Filtering for mesenchymal progenitors and chondrogenic lineages yielded 17 clusters for analysis (Fig. 4D). *LINC02511* was detected primarily in HIC1⁺ mesenchyme^48^, limb mesenchyme, and ChondroPro1 clusters (Fig. 4E), consistent with its expression in progenitor populations in our dataset. PCDH18, COL6A1, and FOXC1 showed similar cluster-specific expression patterns and co-expression with *LINC02511* across the selected clusters (Fig. 4E, Fig. S4G).

Because maxillofacial bones form through intramembranous ossification, the Erickson et al.^47^ craniofacial atlas provided an additional dataset for examining *LINC02511* expression in this pathway. After filtering for skeletal and progenitor cell types (Fig. 4F), LINC02511 was detected in fewer cells but remained present across mesenchymal and skeletal populations. *PCDH18*, *COL6A1*, and *FOXC1* showed overlapping expression patterns (Fig. 4G, Fig. S4H).

In summary, we here report that successful EO priming using OssiGel is associated with the upregulation of non-coding elements, of which *LINC02511* appears as a potential early regulator, predicted to act as a gene expression enhancer. Importantly, our findings were compared to human embryonic atlases of endochondral and intramembranous bone formation, confirming the presence of *LINC02511* predominantly in the EO program of mesenchymal and chondrogenic lineages.

### LINC02511 is an enhancer of the human endochondral ossification program

Since in silico analysis can identify only correlative associations, the functional role of LINC02511 was experimentally investigated. The *LINC02511* expression was first confirmed in cartilage tissues *in vitro* engineered from BM-MSCs. Interestingly, *LINC02511* signal was concentrated in the cartilage ring and perichondrium, as defined by Safranin-O staining (Fig. 5A–B). Quantification of the staining indicated that *LINC02511* was more frequently detected in nuclei in these regions: over 57.7% of nuclei were *LINC02511*-positive, compared with 32% in the resting area (Fig. 5C). However, mean signal intensity did not differ significantly between areas (Fig. 5D).

**Figure 5.**
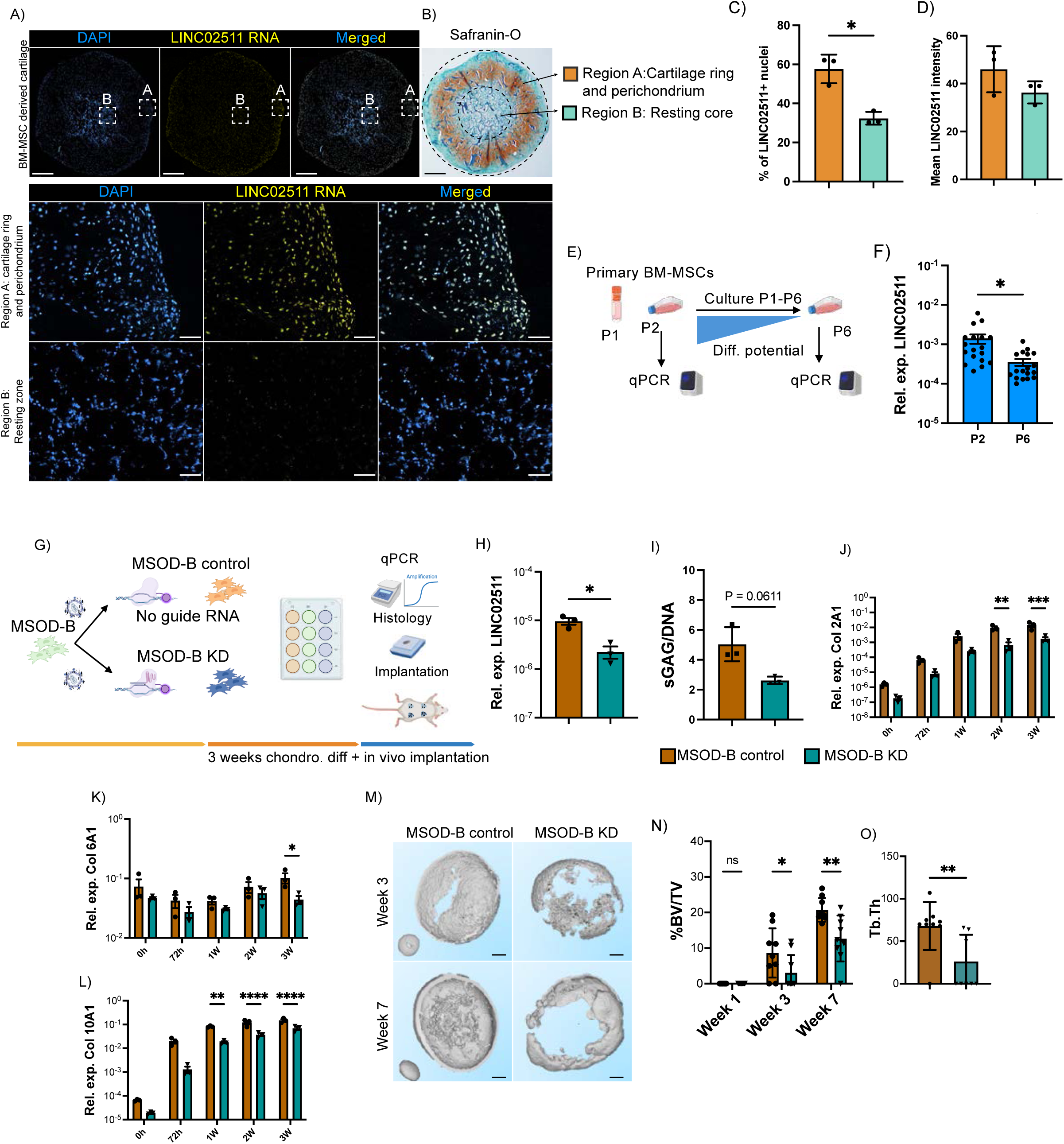
LINC02511 acts as an enhancer in the EO program. (A)LINC02511 RNA fish staining on *in vitro* generated cartilage tissues. Scale bar 1mm (Overview) and 50µm (magnified regions) (B) Safranin-O staining of contiguous section to (A). Scale bar, 1 mm. (C) Image based quantification of % of LINC02511 + nuclei on selected regions. (D) Image based quantification of LINC02511 staining intensity. (E) Experimental scheme for evaluating LINC02511 as a predictor of chondrogenic capacity in primary BM-MSC samples. (F) LINC02511 expression in 15 primary BM-MSC samples at early (P2) versus late (P6) passages. (G) Schematic of the CRISPRi-mediated LINC02511 knockdown design in MSOD-B cells, using two vectors: one encoding dCas9-ZIM3-KRAB with guide RNAs and control with no guides. (H) qPCR validation of *LINC02511* knockdown efficiency. (I) sGAG-to-DNA ratio in chondrogenic pellets from control and LINC02511-knockdown cells. (J-L) *COL2A1* (J), *COL6A1* (K) and *COL10A1* (L) expression during the chondrogenic differentiation time course. (M) μCT reconstruction of ectopic bone formation from control and knockdown cells at 3 and 7 weeks *in vivo.* (N) BV/TV quantification at 1, 3, and 7 weeks post-implantation. (O) Trabecular thickness quantification from control and knockdown cells at 7 weeks *in vivo*. Data are means ± SEM (F, H, J, K, L) or means ± SD (C, D, I, N, O). n=3 BM-MSC derived tissues (C, D), n= 15 BM-MSC samples (F), n=3 differentiation runs (H-L) and n= 9 tissues/condition (N-O). *P < 0.05, **P < 0.01, ****P < 0.0001 by t-test (C, D, F, H, I, O) or two-way ANOVA with multiple comparisons (J, K, L, N).

Ex vivo-expanded BM-MSCs progressively lose their multilineage differentiation capacity with passaging^49^. To determine whether culture-associated aging also affects *LINC02511* expression, we measured its expression in BM-MSCs from 15 donors at early passage (P2) and after further expansion to passage 6 (P6; Fig. 5E). *LINC02511* expression was significantly lower at P6 than at P2, indicating a clear reduction upon prolonged in vitro expansion (Fig. 5F).

Next, the potential role of *LINC02511* in human EO was interrogated by applying CRISPRi to repress its transcription via RNA polymerase blocking, thus turning the gene off without DNA sequence alteration. Because viral transduction can alter the differentiation potential of primary BM-MSCs, we silenced *LINC02511* in MSOD-B cells as a human mesenchymal cell line with stable and well-characterized chondrogenic and EO capacity ^18,50^ (Fig. 5G). A control MSOD-B line lentivirally transduced with the dCas9-ZIM3-KRAB^51^ system alone was generated (MSOD-B control), while the MSOD-B line knockdown for *LINC02511* (MSOD-B KD) resulted from the transduction with four gRNAs targeting a 400 bp window upstream of the *LINC02511* promoter (Fig. 5G, Fig. S5B). The *LINC02511* knockdown efficacy was initially assessed by qPCR of *in vitro* expanded lines, demonstrating a stable 5-fold downregulation of LINC02511 expression in the MSOD-B KD line relative to control (Fig. 5H).

The impact of *LINC02511* knockdown on chondrogenesis was first assessed *in vitro*. Immunostainings of chondrogenic differentiated tissues indicated a reduced type II and type X collagen pattern in the knockdown group (Fig. S5C). In line with these observations, GAG/DNA ratio quantification confirmed a reduction in cartilage formation by the MSOD-KD line (Fig. 5I, Fig. S5D). Temporal analysis of *COL2A1, COL6A1* and *COL10A1* gene expression throughout the *in vitro* time course of differentiation revealed that both MSOD-B control and MSOD-B KD were responsive to the differentiation cues, with a progressive increase in expression (Fig. 5J-L). However, MSOD-B KD exhibited consistent reduction of all three genes at different timepoints, suggesting an impaired differentiation in absence of *LINC02511*.

We next assessed the function of *LINC02511* on endochondral bone formation by implanting MSOD-B control and MSOD-B KD chondrogenic tissues *in vivo*. We performed μCT analysis at 3 and 7 weeks post-implantation (Fig. 5M-O) to monitor calcification. At weeks 3, MSOD-B KD exhibited a reduced pattern of cortical bone formation as compared to the control. This was confirmed at week 7, whereby the MSOD-B control generated a fully formed bone organ with cortical and trabecular structures. In sharp contrast, the cortical shell was incompletely formed, and no trabecular bone could be observed in MSOD-B KD tissues. Those observations were confirmed by quantification of BV/TV (Fig. 5N) and trabecular thickness (Fig. 5O) illustrating the impaired ossification, corroborated by histological analysis (Fig. S5E). Altogether, our data indicate that LINC02511 silencing affects BM-MSCs in vitro chondrogenesis, and significantly impaired bone formation. Together with its nuclear localization, this suggests a key role as enhancer of the human EO process. In line with this, LINC02511 decreases along with BM-MSCs artificial aging performed by *in vitro* passaging.

## Discussion

In this study, we combined a novel tissue engineering technology with single-nucleus multi-omics to provide a comprehensive understanding of epigenetic and transcriptomic changes occurring during the initial steps of human EO. We reveal that successful EO priming is underpinned by chromatin accessibility remodeling at chondrogenic loci and the early emergence of distinct Chondro-progenitors and TWIST1+ MSSCs. We further describe LINC02511 as a novel regulator of chondrogenesis and subsequent EO, functionally demonstrated by CRISPRi-mediated silencing.

Research on human stem cell differentiation remains challenged by poor reproducibility of protocols and pronounced donor-to-donor variability. This particularly applies to BM-MSCs, the primary cell source for cartilage and bone tissue engineering applications, where a minority of donors (∼30%) reliably undergo EO^19,52^. This heterogeneity limits mechanistic studies but also hampers their clinical translation. We here tackled this challenge by exploiting OssiGel, a jellified hypertrophic cartilage matrix engineered by a stable human mesenchymal line. OssiGel performance relies on a combination of osteoinductive factors (BMP-2) and glycosaminoglycans, demonstrated to overcome the limited differentiation capacity of BM-MSCs isolated from leukemic patients^22^. Compared to gold-standard 3D differentiation protocols, OssiGel demonstrated a strong chondro-inductive capacity and enabled reproducible recapitulation of EO across donors. Importantly, this system supported the reconstitution of key mesenchymal lineages within mature bone and bone marrow, indicating preservation of BM-MSC multipotency. Establishing such a controlled and reproducible system provides a foundation to interrogate early EO mechanisms in a human context.

Beyond variability, progress in understanding human EO is further limited by restricted access to relevant biological material motivating the use of a tissue engineering strategy. EO occurs during skeletal development or fracture repair, contexts in which human samples are scarce and difficult to obtain. Consequently, most available BM-MSC datasets derive from healthy, homeostatic bone marrow rather than actively differentiating cells undergoing EO^5,6^. In addition, the rarity of relevant BM-MSC populations further constrains direct analysis. Together, these challenges help explain the scarcity of studies investigating human EO at high resolution^32,53,54^. In this study, we leveraged *in vitro* BM-MSCs expansion, which is known to partially select for clonally dominant mesenchymal populations^55^ reducing heterogeneity while preserving differentiation capacity. Despite these constraints, our model successfully recapitulated EO and its associated cellular lineages. Importantly, our dataset exhibited substantial concordance with recently published embryonic bone formation atlases, further supporting the relevance of our approach.

The establishment of a robust EO model provided a foundation for applying multi-omics approaches to interrogate this process in a human context, aligning with recent paradigms at the convergence of bioengineering, multi-omics, and genetic perturbation^56^. While similar integrative strategies have been applied in other stem cell systems, such as brain and endoderm organoids^57–60^, they remain largely unexplored in the skeletal system. Importantly, despite progresses in organoid and on-chips approaches^56^, EO cannot be fully recapitulated in vitro^61^. To our knowledge, our study provides the first combined single-nucleus transcriptomic and chromatin accessibility mapping of BM-MSC, here applied to the study of EO. Previous epigenetic studies have primarily focused on MSC tissue source-dependent differences, identifying DNA methylation patterns associated with EO potential but limited to bulk analysis^7,62^. Here, our data reveal that chondrogenic differentiation requires extensive chromatin remodeling, characterized by the opening of chondro-specific and osteogenic regulatory regions. This is not obvious, as contrasting with the osteogenic lineage, previously reported to not require a chromatin remodeling step for effective differentiation, as well as mutually exclusive activation of transcription factor networks^5^. These findings indicate that BM-MSCs must actively reconfigure their chromatin landscape to enter the chondrogenic lineage, potentially explaining the higher donor discrepancy in successful chondrogenesis and EO.

Importantly, chromatin remodeling alone was not sufficient to predict successful EO, as accessibility changes were also observed in control Fibrin condition. Instead, successful EO was marked by the emergence of a Chondroprogenitor and TWIST1+ MSSCs populations in transcriptomic data, detected exclusively in the OssiGel condition, highlighting the importance of extrinsic matrix-embedded signals. These findings indicate that successful EO requires the combined establishment of a chondrogenic chromatin landscape leading to early progenitor populations emergence, which can only be capture by multi-omics analysis.

The further characterization of our single nuclei transcriptomic dataset led to the identification of the novel regulator LINC02511, highly upregulated in the OssiGel condition, thus to the EO primed BM-MSCs. Regulation of BM-MSC differentiation has primarily been attributed to master transcription factors and microRNAs ^63–66^. Instead, lncRNAs have shown great biological relevance in other biological settings as in the case of Xist being the prime example of epigenetic inactivation for the X-chromosome^67,68^. LncRNAs have a few distinct mechanisms of action, from modulation of chromatin remodelers, to serving as scaffolding by transcription factors or acting as genetic sponges for mi-RNAs^38,67^. In skeletal system, lncRNAs information remains scarce although recent studies reported the role of lncRNAs ROR^69^ and ROCR^39^ as enhancers of chondrogenic differentiation. No information on the function of LINC02511 has been previously identified, beyond an established link to bone density trait by GWAS analysis^70^.

Our study thus expands the regulatory landscape of EO by identifying LINC02511 as a key modulator of chondrogenesis and bone formation. Functional validation of lowly expressed non-coding elements is challenged by their elusive detection and the scarcity of available methodologies. Due to limited annotation of LINC02511 regulatory regions, we opted for CRISPRi to target a broad promoter region rather than generating complete knockouts. This strategy enabled the rapid generation of an edited MSOD-B cell line, avoiding the challenging of BM-MSC single-clone expansion required for conventional knockout approaches and associated with risks of exhaustion. Although CRISPRi-mediated silencing is partial and reversible, which may attenuate phenotypic effects^51,71^, we identified a clear regulatory role of LINC02511 impacting chondrogenesis and *in vivo* bone formation.

In primary BM-MSCs, reduced LINC02511 expression correlated with declining chondrogenic differentiation potential during extended *in vitro* culture. In silico analysis, together with RNA-FISH imaging notably showing nuclear enrichment of LINC02511 in cartilage regions of BM-MSC-derived tissue, supports a potential enhancer-like role. We postulate that LINC02511 may function in part by promoting expression of COL6A1, which is essential for early EO, as well as BRSK2, a key factor necessary to induce direct reprogramming of placenta derived MSCs toward chondrocytes^72,73^. Future work will help further defining the regulatory function and mechanisms of action of LINC02511, using complete knockout or overexpression systems, studying its expression in larger donor cohorts with clinical metadata, and investigating LINC02511-chromatin interactions, such as ChiRP-seq, although this remains as a big challenge given the low expression of LINC02511.

Focusing on early differentiation stages allowed us to capture the initial EO commitment events, which remain poorly understood yet are critical determinants of downstream outcomes. This approach also provides an opportunity to identify predictive markers of EO potential, a long-standing challenge in the field. Previous efforts have largely relied on surface marker profiling or steady-state transcriptomic signatures, with limited predictive power. Although certain markers have been associated with lineage bias^54,74,75^, their low abundance and variability across donors limit their practical application. Our results suggest that differentiation potential is more accurately encoded within the chromatin landscape and epigenetic state of BM-MSCs rather than surface phenotype alone. While LINC02511 emerges as a key regulator of EO, it is not sufficient as a standalone predictive marker of donor potency. Instead, it likely represents one component of a broader regulatory network involving additional lncRNAs and microRNAs. We therefore hypothesize that combinatorial epigenetic signatures will be required to robustly predict EO capacity, an avenue that warrants further investigation.

In conclusion, our study establishes a robust experimental platform to investigate the cellular and molecular mechanisms governing human skeletal development and repair. It advances understanding of the EO process and highlights the key role of non-coding RNA networks in regulating BM-MSC fate decisions. Considering the importance of BM-MSCs in regenerative medicine, these findings may inform efforts to direct their differentiation for therapeutic applications.

### Limitations of the study

This study has several limitations. First, our analyses focus on a single early timepoint, capturing initial events but not the full temporal dynamics of epigenetic and transcriptional regulation during EO. Longitudinal studies, combined with additional approaches such as proteomics, spatial transcriptomics, or chromatin interaction assays, would provide a more comprehensive view of regulatory mechanisms. Second, although our tissue engineering–based model enables controlled and reproducible investigation of human EO, it relies on ectopic implantation in immunodeficient mice and may not fully recapitulate the native human physiological environment. While supported by concordance with human skeletal atlases, differences with *in vivo* human processes cannot be excluded. Of note, despite significantly enhancing chondrogenic potential consistently, OssiGel does not completely overcome donor variability in cartilage formation.

## Resource availability

The single cell multiomic datasets generated in this study have been made available in the Gene Expression Omnibus (GEO) under accession number GSE339419.

The code and resources for the construction of the singularity environment analysis and analysis scripts are available at GitHub under the repository: https://github.com/davhg96/ChromAcc_EO

A complete singularity image can be shared upon request.

## Acknowledgements

We thank the Lund Stem Cell Center FACS Core Facility for assistance with flow cytometry and cell sorting experiments. We are grateful to the Lund Stem Cell Center Imaging Facility and the Lund Bioimaging Centre (LBIC) for access imaging equipment and support. The authors would like to acknowledge Clinical Genomics Lund, SciLifeLab and Center for Translational Genomics (CTG), Lund University, for providing expertise and service with sequencing. We would like to thank the Centre for Comparative Medicine (CCM) for their expertise and support with animal handling and experimentation. We thank Dr. Luis Quintino for valuable assistance with CRISPR design and technical guidance. We would like to thank Dr. Aurélie Baudet for her support and feedback on the manuscript. We also would like to thank Dr. Camille Sauter for her support and expertise on animal experiments and Dr. Dimitra Zacharaki for the isolation and culture of BM-MSC samples.

This project has been funded by the European Research Council (Starting grant #948588 to P.E.B), from the Swedish Research Council (2019-01864 to P.E.B.), Cancerfonden to P.E.B, Sten K Johnsons Stiftelse and the IngaBritt och Arne Lundbergs Forskningsstiftelse (to P.E.B).

## Author contributions

Conceptualization, D.H.G., and P.E.B.; Methodology, D.H.G., A.G.G., and P.E.B.; Formal Analysis, D.H.G.; Investigation, D.H.G., A.G.G., F.W., S.G.A., S.B., A.G., P.C.T.N., and S.M.; Data Curation, D.H.G.; Visualization, D.H.G.; Writing – Original Draft, D.H.G. and P.E.B.; Writing – Review & Editing, D.H.G., A.G.G., F.W., S.G.A., S.B., A.G., P.C.T.N., S.M., C.B., A.B., and P.E.B.; Supervision, P.E.B.; Project Administration, D.H.G. and P.E.B.; Funding Acquisition, D.H.G. and P.E.B.

## Supplemental information titles and legends

Supplemental table 1. BM-MSC donor demographics

Supplemental table 2. LongTarget results for LINC02511

Supplemental table 3. catRAPID RNA-Protein interactions for LINC02511

**Supplementary Figure S1.**
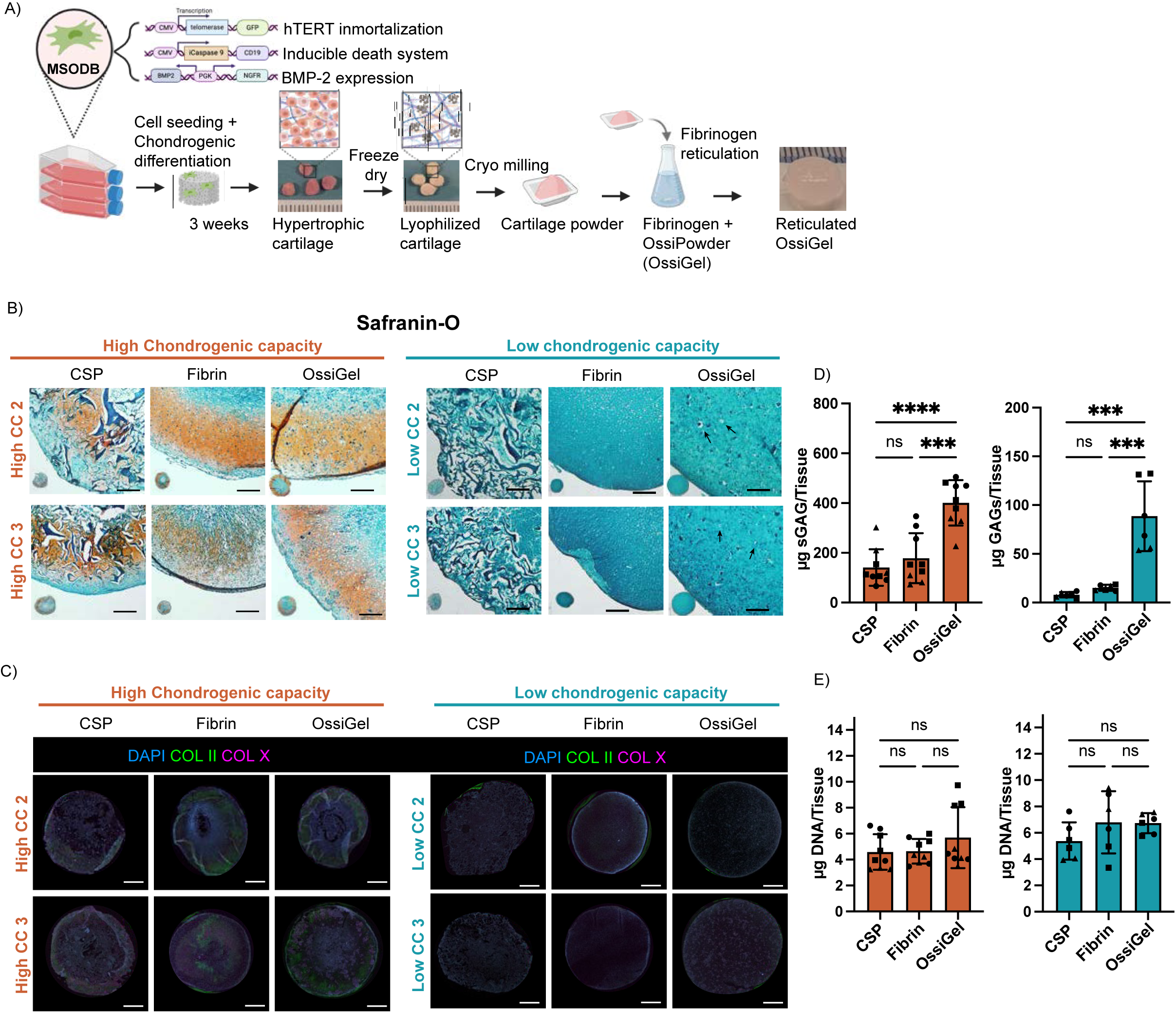
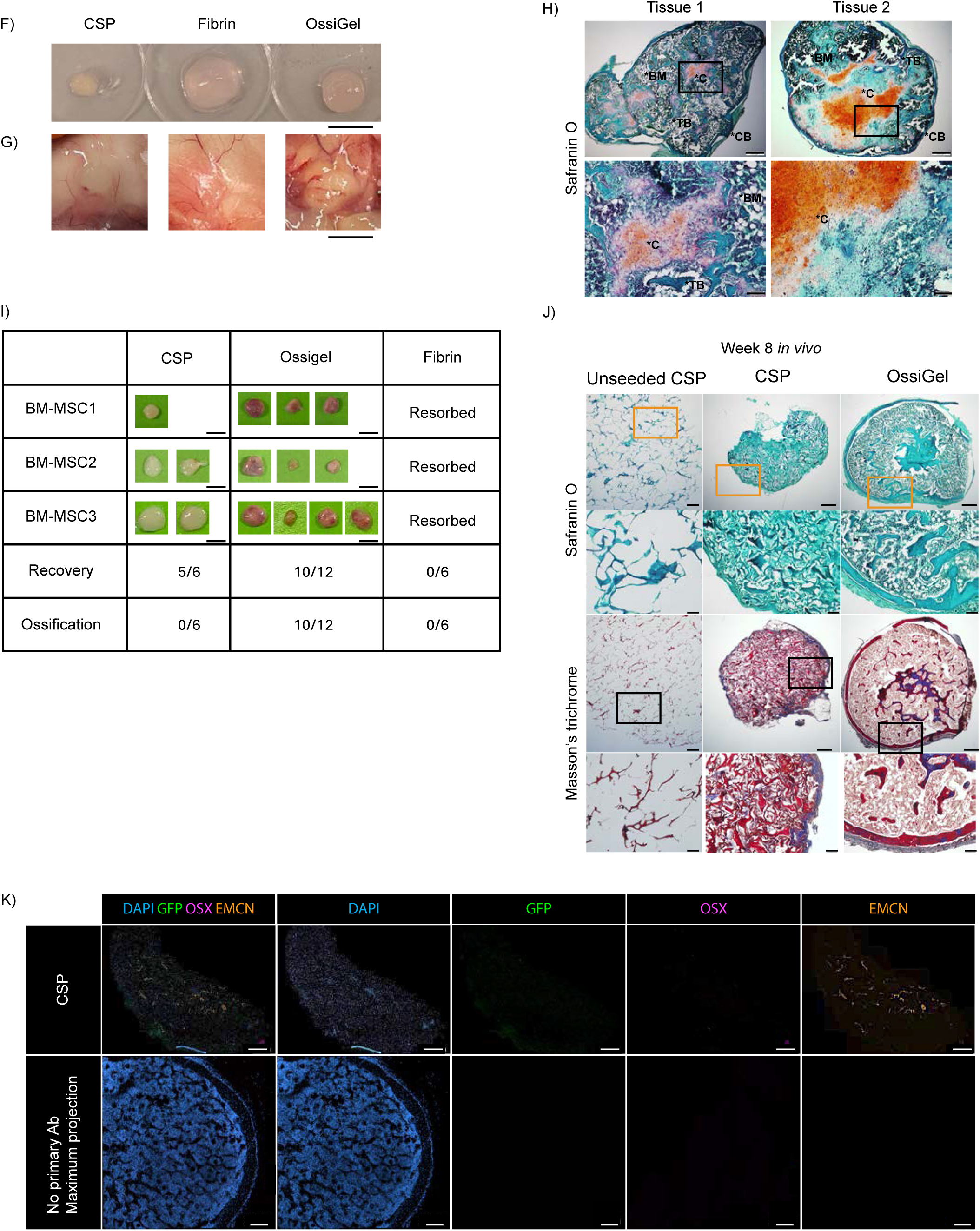
OssiGel confers superior chondrogenesis priming of human bone marrow mesenchymal stem/stromal cells and robust recapitulation of endochondral ossification. (A) Schematic of OssiGel production. (B) Safranin-O staining of biological replicates from high-CC (left) and low-CC (right) samples in CSP, fibrin, and OssiGel. Arrows indicate chondrocyte lacunae. Scale bars, 500µm. (C) Immunofluorescence for type II and type X collagens in biological replicates from high-CC (left) and low-CC (right) samples in CSP, fibrin, and OssiGel. Scale bars, 1 mm. (D and E) sGAG (D) and DNA (E) quantification after 3 weeks of chondrogenic differentiation in CSP, fibrin, or OssiGel scaffolds. (F) Macroscopic images of *in vitro*–generated tissues prior to implantation. Scale bar, 10 mm. (G) Macroscopic view of implantation sites for CSP, fibrin, and OssiGel after 3 days *in vivo*. Scale bar, 10 mm. (H)Safranin-O staining of OssiGel derived ossicles at 6-weeks *in vivo* showing cartilage (C), cortical bone (CB), bone marrow (BM), and trabecular bone (TB). Scale bars, 500 μm (overview) and 100 μm (magnified regions). (I) Macroscopic images of 8-week explants showing bone and marrow formation in OssiGel conditions, with recovery and ossification rates shown per donor. Scale bar, 5 mm. (J) Safranin-O and Masson’s trichrome staining of CSP and OssiGel explants at 8 weeks *in vivo*. Scale bars, 500 μm (overview) and 100 μm (magnified regions). (K) Immunofluorescence of CSP explants (top) and no-primary-antibody control (bottom) for DAPI, GFP, OSX, and EMCN, showing lack of hBM-MSC persistence in collagen scaffolds. Scale bars, 500 μm. Data are means ± SD; n = 3 biological replicates per group. *P < 0.05, **P < 0.01, ***P < 0.001, ****P < 0.0001 by two-way ANOVA with multiple comparisons (D, E).

**Supplementary Figure S2.**
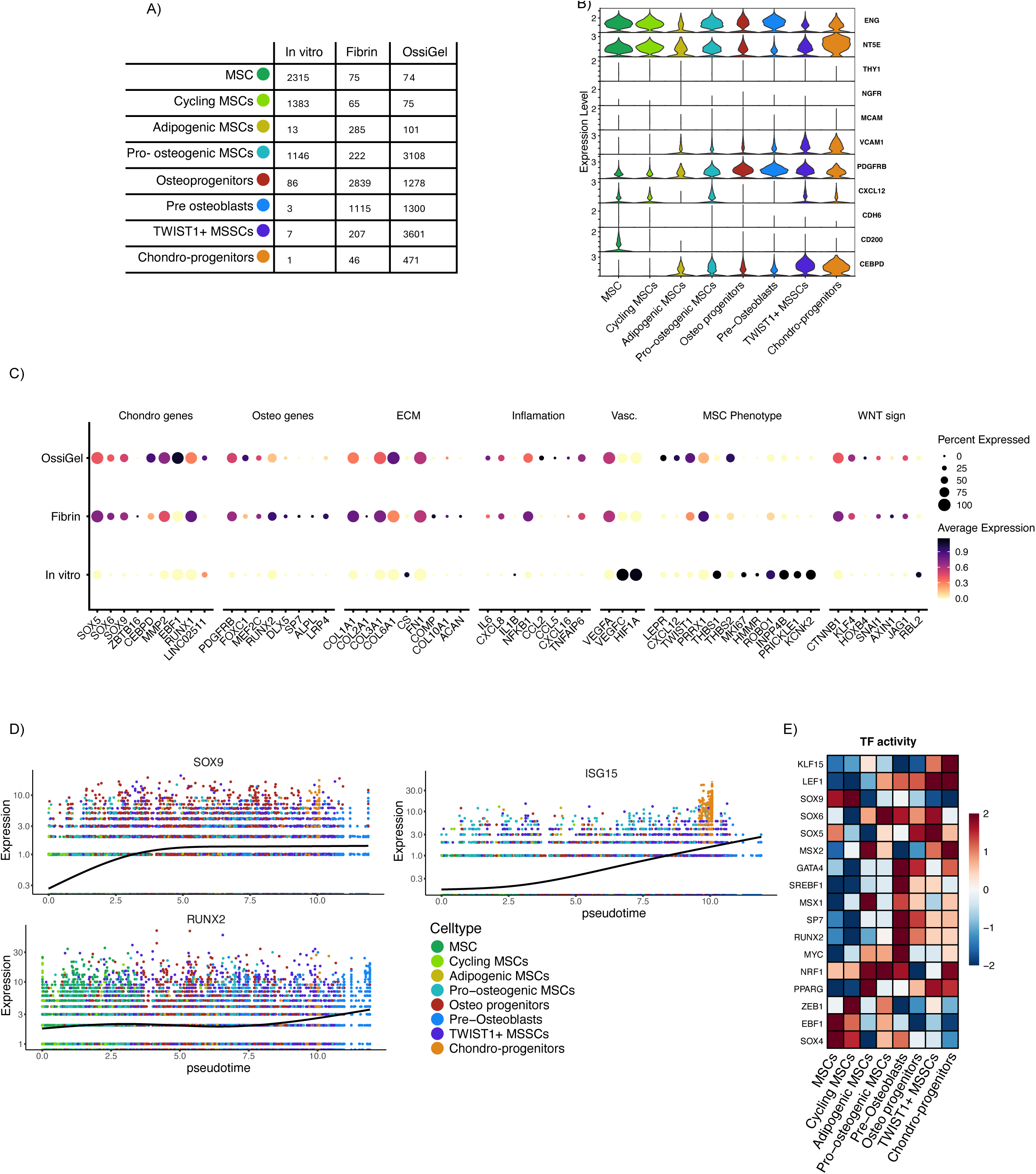
Early emergence of Chondro-progenitors and TWIST1*+* mesenchymal skeletal stem cells is a hallmark of endochondral ossification priming. (A) Cell counts per cluster across experimental conditions. (B) Violin plots of selected stromal markers. (C) Dot plot of average scaled expression (color) and fraction of expressing cells (dot size) for genes grouped by functional category (chondrogenic, osteogenic markers, extracellular matrix, inflammation, vasculogenesis, MSC phenotype markers and WNT signaling) across conditions. (D) Pseudotime ordered expression of *SOX9, ISG15* and *RUNX2*. (E) Scaled transcriptional activity of selected transcription factors across clusters.

**Supplementary Figure S3.**
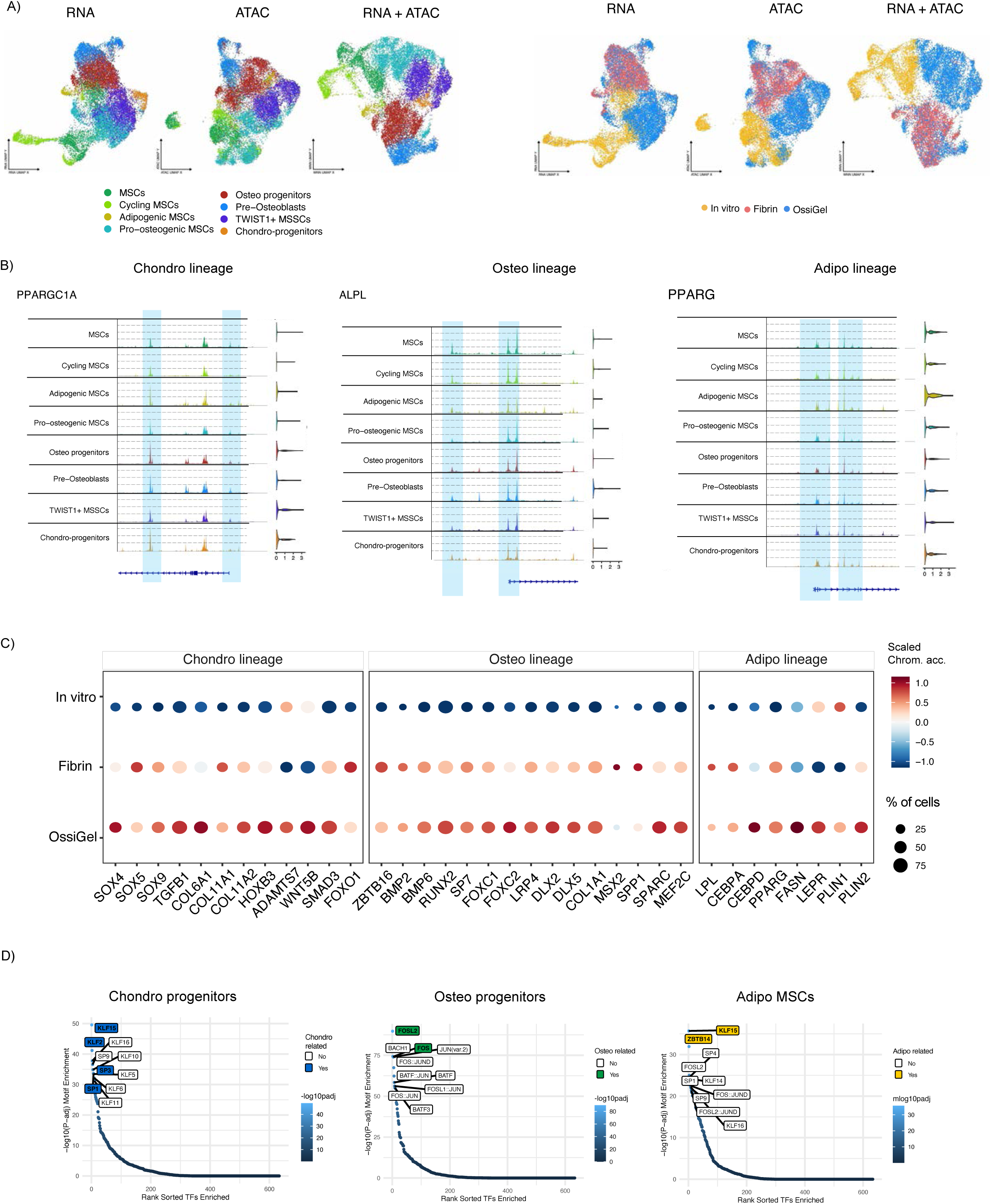
Endochondral ossification is associated with early and distinct chondrogenic chromatin remodeling signatures. (A) UMAP embeddings of RNA, ATAC, and weighted RNA + ATAC embedding, colored by cell population (left) and sample (right). (B) Genome track plots of *PPARGC1A*, *ALPL*, and *PPARG* loci with corresponding gene expression levels, highlighted are selected regions of interest. (C) Dot plot of average scaled chromatin accessibility (color) and fraction of accessible cells (dot size) for selected marker genes by condition. (D) Ranked transcription factor motif enrichment in Chondro-progenitor (left), Pre-Osteoblast (middle), and Adipogenic MSCs (right) clusters

**Supplementary Figure 4.**
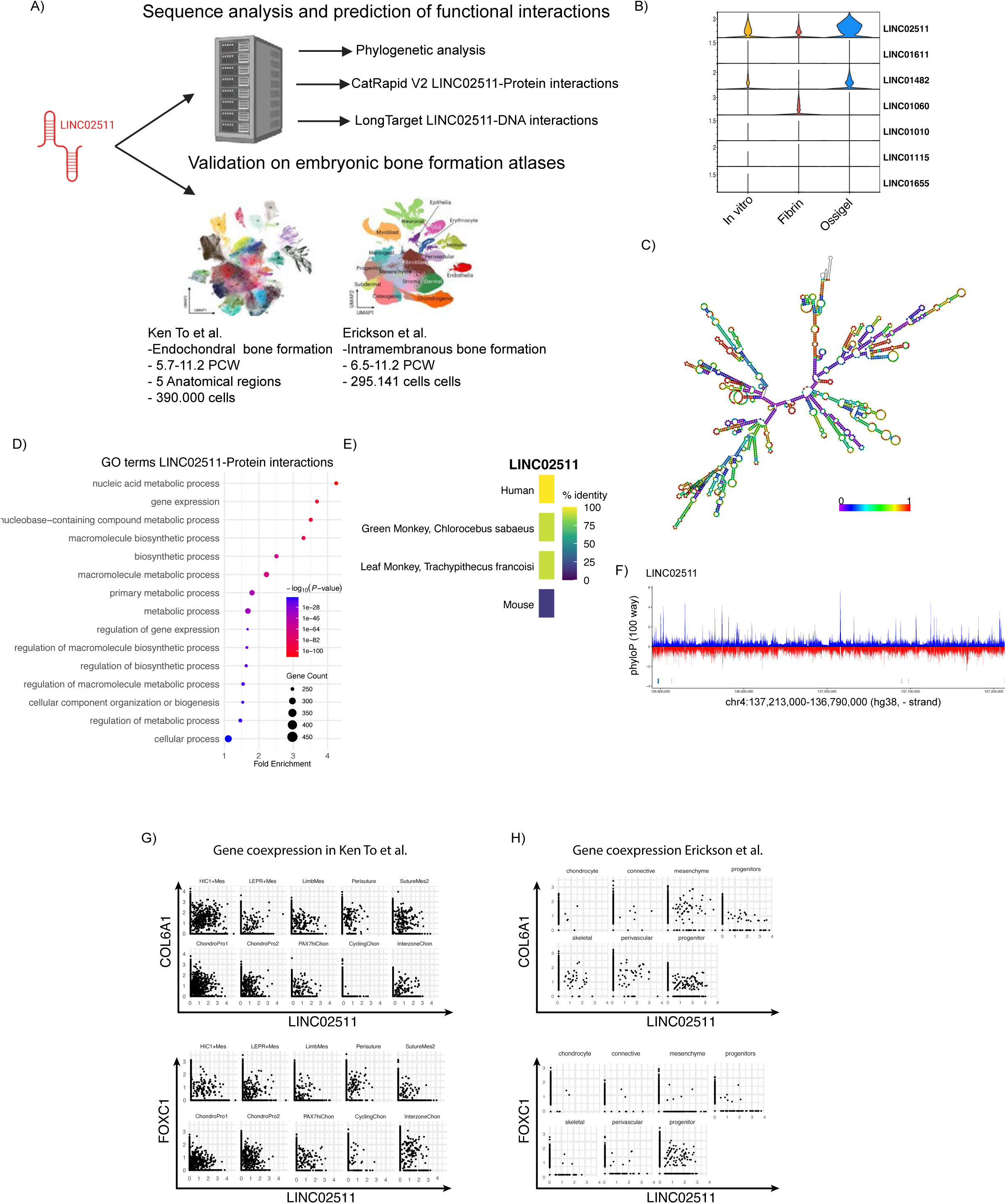
LINC02511 expression is a feature of human embryonic bone development. (A) Schematic representation of computational analysis performed on *LINC02511* for the prediction of RNA-genome and RNA-protein interactions and expression validation in embryonic bone development atlas. (B)Violin plot of differentially expressed long non-coding RNAs in our dataset by condition. (C) Predicted secondary structure of *LINC02511* (D) Gene ontology enrichment Dot plot of proteins with predicted interactions with LINC02511. (E) Percentage of homology of *LINC02511* across species. (F) PhyloP100 analysis of *LINC02511*. (G-H) Co-expression of *LINC02511* and its putative target gene *COL6A1* and *FOXC1* across selected cell types of Ken To et. al, Nature, 2024 (G) and Erickson et al., Nature Genetics 2026 (H)

**Supplementary Figure S5.**
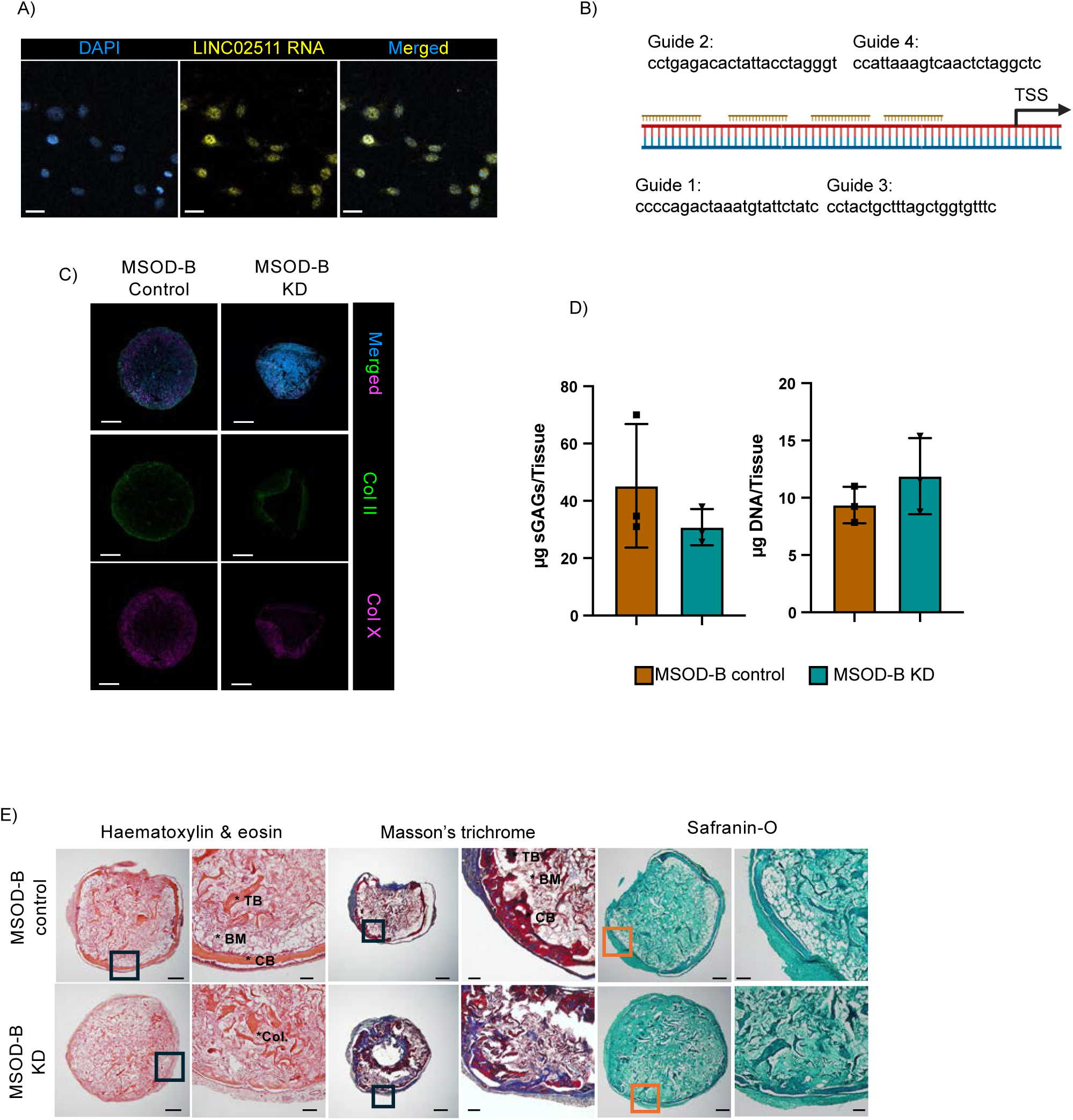
LINC02511 acts as an enhancer in the EO program. (A) RNA fluorescence in situ hybridization (FISH) for *LINC02511* in cultured MSOD-B cells. Scale bar, 10 μm. (B) Sequences and genomic location of CRISPRi guide RNAs targeting the LINC02511 transcription start site (TSS). (C) Representative immunofluorescence for type II and type X collagens in chondrogenic pellets of LINC02511 KD and control cell lines. (D) sGAG (left) and DNA (right) quantification in MSOD-B control versus LINC02511-knockdown tissues. (E) Histological analysis (H&E, Masson’s trichrome, Safranin-O) of 7-week implanted cartilage tissues derived from MSOD-B KD and control cell lines, showing cortical bone (CB), trabecular bone (TB), bone marrow (BM), and unremodelled collagen sponge (CSP).

## Methods

### Healthy donor BM samples

Bone marrow aspirates were obtained from willing healthy donors under informed written consent in accordance with the Declaration of Helsinki under ethical approval: 2021-04046.

BM aspirates were filtered and diluted 1:2 in PBS. Mononucleated cells were isolated by density gradient centrifugation using LMS 1077 Lymphocyte (PAA Labs, Austria). Mononucleated cells were seeded at a density of 0.2×10^6^/cm^2^ for MSC isolation by plastic adherence.

### BM-MSC Isolation and culture

BM-MSCs were seeded at a density of 3200 cells/cm² and cultured until reaching 80-90% confluency in alpha-minimum essential medium (α-MEM) supplemented with 10% fetal bovine serum (FBS), 1% HEPES (1 M), 1% sodium pyruvate (100 mM), 1% penicillin-streptomycin-glutamine (100×) solution (all from Gibco), and 5 ng/ml fibroblast growth factor 2 (FGF-2; R&D Systems) in a humidified 37°C/5% CO₂ incubator. The medium was changed twice weekly. Upon reaching confluency, cells were detached using 0.05% trypsin-EDTA (Gibco) and expanded for subsequent experiments. All primary BM-MSC were used at P<5, unless otherwise stated, while MSOD-B were used at P<22.

### Chondrogenic differentiation of BM-MSCs

Two million BM-MSCs were seeded into 6 mm diameter bovine type I collagen sponges (Ultrafoam 1050050) and cultured for 3 weeks in chondrogenic medium consisting of Dulbecco’s modified Eagle’s medium supplemented with 1% penicillin-streptomycin-glutamine (Gibco), 1% HEPES (1 M) (Gibco), 1% sodium pyruvate (100 mM) (Gibco), 1% ITS (100×) (insulin, transferrin, and selenium) (Gibco), 0.47 mg/ml linoleic acid (Sigma-Aldrich), 0.12% bovine serum albumin (Sigma-Aldrich), 0.1 mM ascorbic acid (Sigma-Aldrich), 10⁻⁷ M dexamethasone (Sigma-Aldrich), and 10 ng/ml transforming growth factor-β3 (Novartis).

### OssiGel matrix production

MSOD-B cells at passage <22 were seeded at 3200 cells/cm² and cultured until reaching confluency. Cells were then detached and seeded into 6 mm diameter collagen sponges and cultured in chondrogenic medium for 3 weeks as previously described. MSOD-B-derived cartilage tissues were snap-frozen in liquid nitrogen and freeze-dried overnight. Resulting dry pellets were subsequently ground on a cryomill (Retsch).

### GAG/DNA quantification

Following chondrogenic differentiation, tissue samples were digested in proteinase K overnight at 60°C. Sulfated glycosaminoglycans were quantified from the digested samples using the Blyscan assay (BioColor Life Science Assays #B1000) according to the manufacturer’s instructions. DNA was quantified from the proteinase K lysate using the CyQUANT Cell Proliferation Assay (Fisher Scientific #C35006).

### Histology of paraffin tissues

After chondrogenic differentiation, tissues were fixed in 4% formaldehyde overnight at 4°C. Fixed tissues were dehydrated through a series of ethanol immersion baths: 70% EtOH for 40 min, 95% EtOH for 40 min, 99.5% EtOH for 40 min, 99.5% EtOH/xylene (1:1 v/v) for 10 min, and xylene for 40 min. Dehydrated tissues were incubated in infiltration paraffin (Merck #P2558) at 60°C overnight. Tissue blocks were sectioned at 10 μm thickness, mounted on SuperFrost Plus slides (VWR 95057985), and incubated at 40°C overnight to dry.

#### Safranin-O staining

Dried 10 μm tissue sections were deparaffinized and rehydrated through xylene-to-ethanol baths and stained with Mayer’s hematoxylin (Merck MHS80), 0.01% Fast Green (Fisher Scientific A16520.22), and 0.001% Safranin-O (Fisher Scientific B21674.18). Tissues were then cleared and dehydrated through graded ethanol baths and xylene.

#### Modified Bern score (MBS)

Representative Safranin-O staining images for each sample were blindly evaluated by three independent researchers and scores ranging 0-3 were assigned for cell morphology and Safranin-O staining intensity respectively. Scores are added for each category and average score across researchers is obtained for each tissue.

#### Hematoxylin and eosin

Dried 10 μm tissue sections were deparaffinized and rehydrated through xylene-to-ethanol baths and stained with Mayer’s hematoxylin (Merck MHS80), followed by 5% eosin Y (Fisher Scientific 201930250) at pH 4.5 for 5 minutes. Tissues were then cleared and dehydrated through graded ethanol baths and xylene.

#### Mason’s trichrome staining

Masson’s trichrome staining was performed on 10 μm tissue sections according to the manufacturer’s instructions (Sigma-Aldrich HT15).

### Ectopic implantation in mouse

NSG or nude female mice (2 to 4 months old) were obtained from Charles River Laboratories. Each mouse was implanted 4 *in vitro* engineered tissues. All mouse experiments and animal care were performed in compliance with the Lund University Animal Ethical Committee (15485-18 and 1901219). Tissue implantations were done subcutaneously in the back of the mice.

### *In vivo* tissue digestion for SN Multiomics

Tissues selected for multiomics sequencing were explanted from the mice and digested in Nattokinase (MedChemExpress #HY-P2373) as described ^25^.

Following isolation, cells were assessed for viability and lysed for nuclei isolation according to the 10X Genomics Demonstrated Protocol CG000365-Rev C. Library construction and sequencing were performed by the Centre for Translational Genomics (CTG Lund) using a 10X Multiome kit. Sequencing was performed on the NovaSeq X Series platform. Reference mapping and feature counting were performed CellRanger-ARC.

### Single nuclei multiomics data analysis

Downstream data analysis was performed following standard procedures and best practices using Seurat v5.0 and Signac v1.14.0. The complete software suite can be found on the GitHub code repository.

### Immunofluorescence staining, imaging and analysis

A complete list of antibodies used is provided in the Supplementary Materials. Immunofluorescence staining of formalin-fixed paraffin-embedded (FFPE) sections was performed on 10 μm thick sections as previously described^767572747576^.

*In vivo* implanted tissues were fixed in 4% paraformaldehyde for 24 hours. Following fixation, tissues were embedded in 4% agarose and sectioned at 100 μm thickness using a vibratome (Campden Instruments). Downstream staining and imaging were performed as previously described^18^.

Images were acquired on a Leica Stellaris 5 with five HyD detectors, and 2 lasers (405mm blue diode and a WLL. Using type G oil (RU 1.45) and a 20X HC PL APO lens with a NA of 0.75. Scans were acquired at 400-600Hz with a z-step of 3-5µm depending on tissue thickness and a tile resolution of 1024×1024 pixels.

Acquired images were processed downstream on IMARIS V17.7 or QuPath V0.7. In selected cases images have been cropped to approximate the region corresponding to the human tissue.

### LINC02511 HCR imaging

HCR imaging was performed following V3 protocol (molecular instruments) for FFPE tissues or cells on a slide respectively according to manufacturer’s instructions. LINC02511 probes were generated using a publicly available probe generator previously described^77^ with a B2 adapter and B2-AF546 amplifier.

### CRISPRi

Guide RNAs (gRNAs) targeting the LINC02511 promoter region were designed using the CHOPCHOP CRISPR toolbox.^78^. Four different gRNAs evenly distributed across the promoter region were selected and cloned into the pHR-UCOE-SFFV-dCas9-mCherry-ZIM3-KRAB vector (Addgene plasmid #154473) ^51^. Control vector (without gRNAs) and knockdown vectors were used to generate lentiviral libraries at the Gene and Cell Therapy Core Facility at Lund University. Following transduction, MSOD-B cells were sorted by fluorescence-activated cell sorting (FACS) based on mCherry reporter expression and subsequently expanded.

### qPCR

Total RNA was isolated from cells using the RNeasy Mini Kit and from chondrogenic differentiated tissues using the RNeasy Fibrous Tissue Mini Kit (Qiagen). One to two micrograms of RNA were reverse-transcribed into cDNA using SuperScript IV VILO (Fisher Scientific #11-754-050). TaqMan-compatible probes for LINC02511 were designed using Primer3^79^ and synthesized by Fisher Scientific. qPCR analysis was performed on a Bio-Rad CFX Opus 384 platform using Fisher Scientific TaqMan qPCR reagents and previously reported primers. Relative expression was calculated using the 2^⁻ΔCt^ method against GAPDH.

